# The murine transcriptome reveals global aging nodes with organ-specific phase and amplitude

**DOI:** 10.1101/662254

**Authors:** Nicholas Schaum, Benoit Lehallier, Oliver Hahn, Shayan Hosseinzadeh, Song E. Lee, Rene Sit, Davis P. Lee, Patricia Morán Losada, Macy E. Zardeneta, Róbert Pálovics, Tobias Fehlmann, James Webber, Aaron McGeever, Hui Zhang, Daniela Berdnik, Weilun Tan, Alexander Zee, Michelle Tan, The Tabula Muris Consortium, Angela Pisco, Jim Karkanias, Norma F. Neff, Andreas Keller, Spyros Darmanis, Stephen R. Quake, Tony Wyss-Coray

## Abstract

Aging is the single greatest cause of disease and death worldwide, and so understanding the associated processes could vastly improve quality of life. While the field has identified major categories of aging damage such as altered intercellular communication, loss of proteostasis, and eroded mitochondrial function^1^, these deleterious processes interact with extraordinary complexity within and between organs. Yet, a comprehensive analysis of aging dynamics organism-wide is lacking. Here we performed RNA-sequencing of 17 organs and plasma proteomics at 10 ages across the mouse lifespan. We uncover previously unknown linear and non-linear expression shifts during aging, which cluster in strikingly consistent trajectory groups with coherent biological functions, including extracellular matrix regulation, unfolded protein binding, mitochondrial function, and inflammatory and immune response. Remarkably, these gene sets are expressed similarly across tissues, differing merely in age of onset and amplitude. Especially pronounced is widespread immune cell activation, detectable first in white adipose depots in middle age. Single-cell RNA-sequencing confirms the accumulation of adipose T and B cells, including immunoglobulin J-expressing plasma cells, which also accrue concurrently across diverse organs. Finally, we show how expression shifts in distinct tissues are highly correlated with corresponding protein levels in plasma, thus potentially contributing to aging of the systemic circulation. Together, these data demonstrate a similar yet asynchronous inter- and intra-organ progression of aging, thereby providing a foundation to track systemic sources of declining health at old age.

---

To uncover aging dynamics organism-wide, we measured plasma proteins and sequenced RNA from 17 organ types isolated from C57BL/6JN males (n=4, aged 1, 3, 6, 9, 12, 15, 18, 21, 24, 27 months) and females (n=2, ages 1, 3, 6, 9, 12, 15, 18, 21 months) (Figure 1A,B). We isolated all 17 organs from each mouse, including bone (both femurs and tibiae), brain (hemibrain), brown adipose tissue (BAT, interscapular depot), gonadal adipose tissue (GAT, inguinal depot), heart, kidney, limb muscle (*tibialis anterior*), liver, lung, marrow, mesenteric adipose tissue (MAT), pancreas, skin, small intestine (duodenum), spleen, subcutaneous adipose tissue (SCAT, posterior depot), and white blood cells (WBCs, buffy coat). Raw data are available from GEO (GSE132040), and an interactive data browser is available at https://twc-stanford.shinyapps.io/maca/.

**Figure 1.**
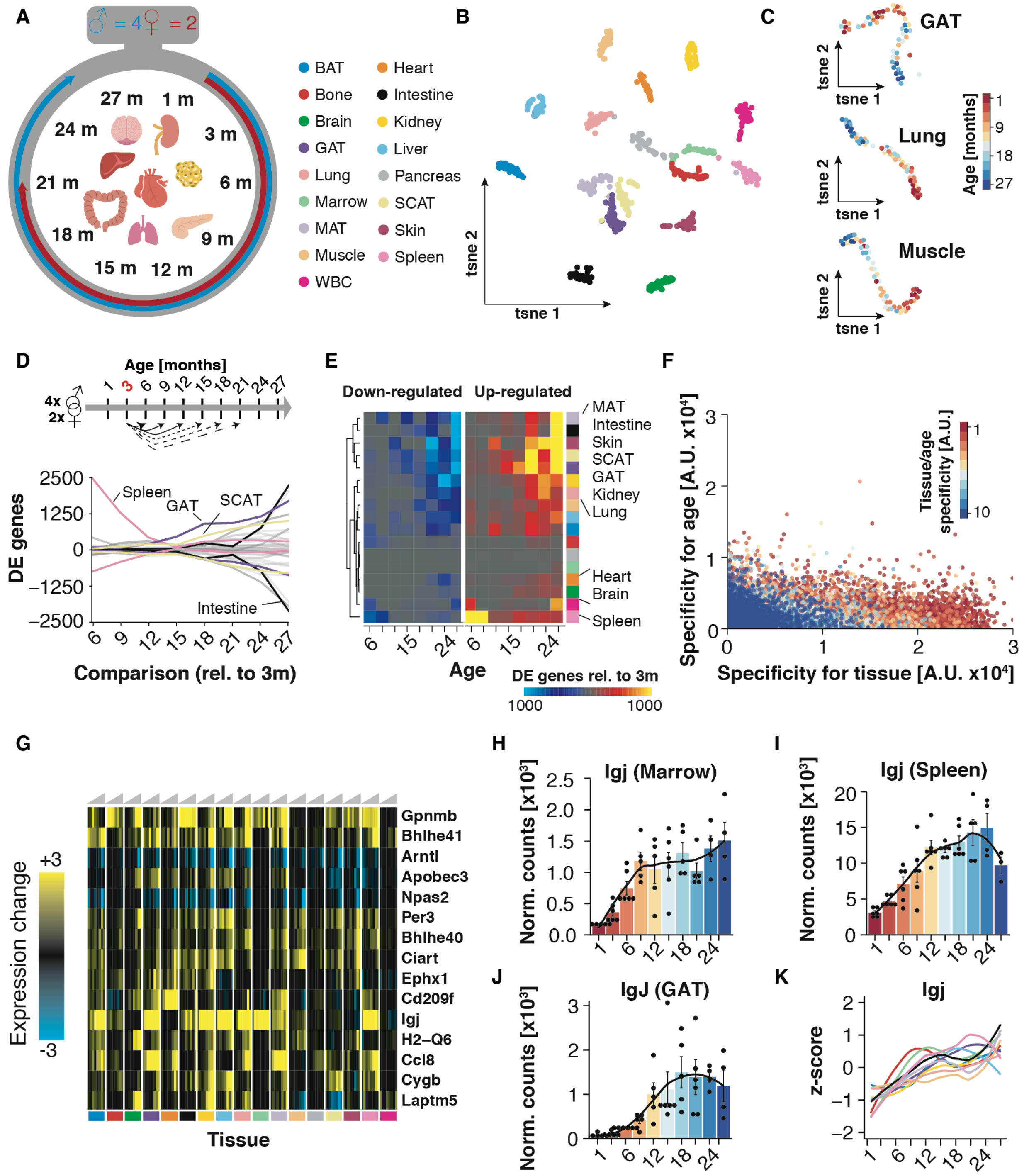
Pairwise differential expression across organs. (A) Experiment outline. 17 organ types were collected from males 1-27mos old (n=4 at each age) and females 1-21mos old (n=2 at each age). (B) t-SNE visualization of all samples. (C) t-SNE visualization of GAT, lung and muscle, colored by age. (D) Smoothed lineplot displaying the number of differentially expressed genes for pairwise comparisons with a 3mo reference, with each tissue being represented by a pair of lines. Positive (negative) values represent up-regulated (down-regulated) genes. (E) Heatmap representation of (D). (F) Scatterplot displaying gene-wise enrichment scores for tissue, age or pair of tissue/age. (G) Tissue-wise expression changes with age (column-wise from left to right) for the top 15 genes exhibiting shifts in most tissues. (H-J) IgJ (Jchain) mRNA expression (RNA-seq) in (H) Marrow, (I) Spleen and (J) GAT. LOESS regression is indicated by black line. (K) Z-transformed, smoothed gene expression trajectory of IgJ, colored by tissue. Means ± SEM.

Aging instigates functional decline across organs, disrupting intricate crosstalk essential for maintaining healthy organismal processes. While individual organs segregate by age (Figure 1C), we lack a basic comparative understanding of aging between these organs, including differences in the onset and rate of aging. We therefore performed pairwise differential expression to determine when differentially expressed genes (DEGs) arise and if they persist with advancing age. While we find few DEGs between neighboring ages, relative to 3-month-old young adults, the number of DEGs increases dramatically in most organs, suggesting progressive, gradual changes that reach detectability only after sufficient time (Figure 1D,E). Some organs, like pancreas and marrow, appear relatively refractory to aging gene expression changes, perhaps explaining the relatively small proportion of global variance due to aging in the dataset (Extended Data Figure 1A). While core aging profiles are maintained relative to 6-month-old mice, DEGs increase greatly with a 1-month-old reference, exemplifying strong developmental phenotypes (Extended Data Figure 2A-C). The clear exception is the spleen, which displays large quantities of DEGs during early life. Notably, SCAT and GAT DEGs arise in mid-life prior to other organs (Extended Data Figure 2D). These may be connected to known changes in adipose composition, especially regarding immune cell infiltration^2^. Some organs, such as mesenteric fat and the small intestine to which it is attached, undergo acute and drastic gene expression changes only late in life. To independently confirm our observations, we generated self-organizing maps for each organ, which allows visualization of correlated gene nodes (Extended Data Figure 3A)^3^. Not only do global aging nodes emerge, but white adipose tissues again exhibit strong aging profiles, with visceral GAT and MAT highly similar. Sex also influences organ function, leading to divergent aging and disease outcomes in humans^4^. We observe prominent sex effects in GAT, SCAT, liver, and kidney, possibly connected with known differences in fat storage, sex hormone regulation, and renal hemodynamics (Extended Data Figure 1B, Extended Data Figure 3B)^5,6,7^.

We next asked if early life DEGs persist with advancing age, or if they give way to new DEGs at older ages. In bone for example, early life expression of ossification genes decreases as bone formation completes, and these genes are highly correlated with late-life DEGs that increase in expression, typifying age-related bone loss (Extended Data Table 1). Overall, most organs show correlation between early and late DEGs, exemplified for nearly every pairwise comparison in GAT, liver, kidney, and heart (Extended Data Table 1). Indeed, few genes are unique for any individual age, with organ-specificity outweighing age-specificity (Figure 1F). Differential expression common between aging organs is especially interesting, as ubiquitous aging pathways may present novel therapeutic opportunities. When we isolated genes most commonly differentially expressed across organs, we found strong enrichment for immune response pathways (Figure 1G, Extended Data Figure 2E). Interestingly, the plasma B cell marker immunoglobulin J (Igj/Jchain) demonstrates a persistent increase throughout life in 11 of 17 organs (Figure 1G-K). Circadian clock genes Bhlhe40/41, Arntl, Npas2, Per3, Ciart, and Dbp also debut among top DEGs (Extended Data Figure 2F). Age-related circadian disruption is well known, but these data perhaps highlight the underappreciated organism-wide role for circadian rhythms in declining health. In fact, a malfunctioning circadian clock appears to contribute to metabolic and inflammatory disorders, and shortened lifespan^8^.

Pairwise comparisons are inherently limited, and our data allow interrogation of gene expression dynamics with high temporal resolution across the lifespan. We first searched for trajectories with common behavior between organs to reveal organism-wide processes. We calculated the average trajectory for each gene across all 17 organs, and clustering those averaged trajectories, revealing functional enrichment for aging hallmarks such as elevated inflammation, mitochondrial dysfunction, and loss of proteostasis (Figure 2A, Extended Data Table 2). Notably, these hallmarks undergo distinct dynamic patterns. For example, cluster 3 declines linearly across the lifespan and is strongly enriched for mitochondrial genes, whereas cluster 7 demonstrates a sharp decline of heat shock proteins important for protein folding, but only beginning at 12 months of age. This is in contrast to cluster 8 extracellular matrix genes which decline rapidly until 6 months, from when a more gradual decline prevails. Immune response pathways feature in clusters 4 and 6. Cluster 4 genes like beta-2 microglobulin (B2m) and Igj increase steadily throughout life. On the other hand, cluster 6 immune genes like Cd74 and complement C1q experience a non-linear increase featured by a plateau between 9 and 15 months.

**Figure 2.**
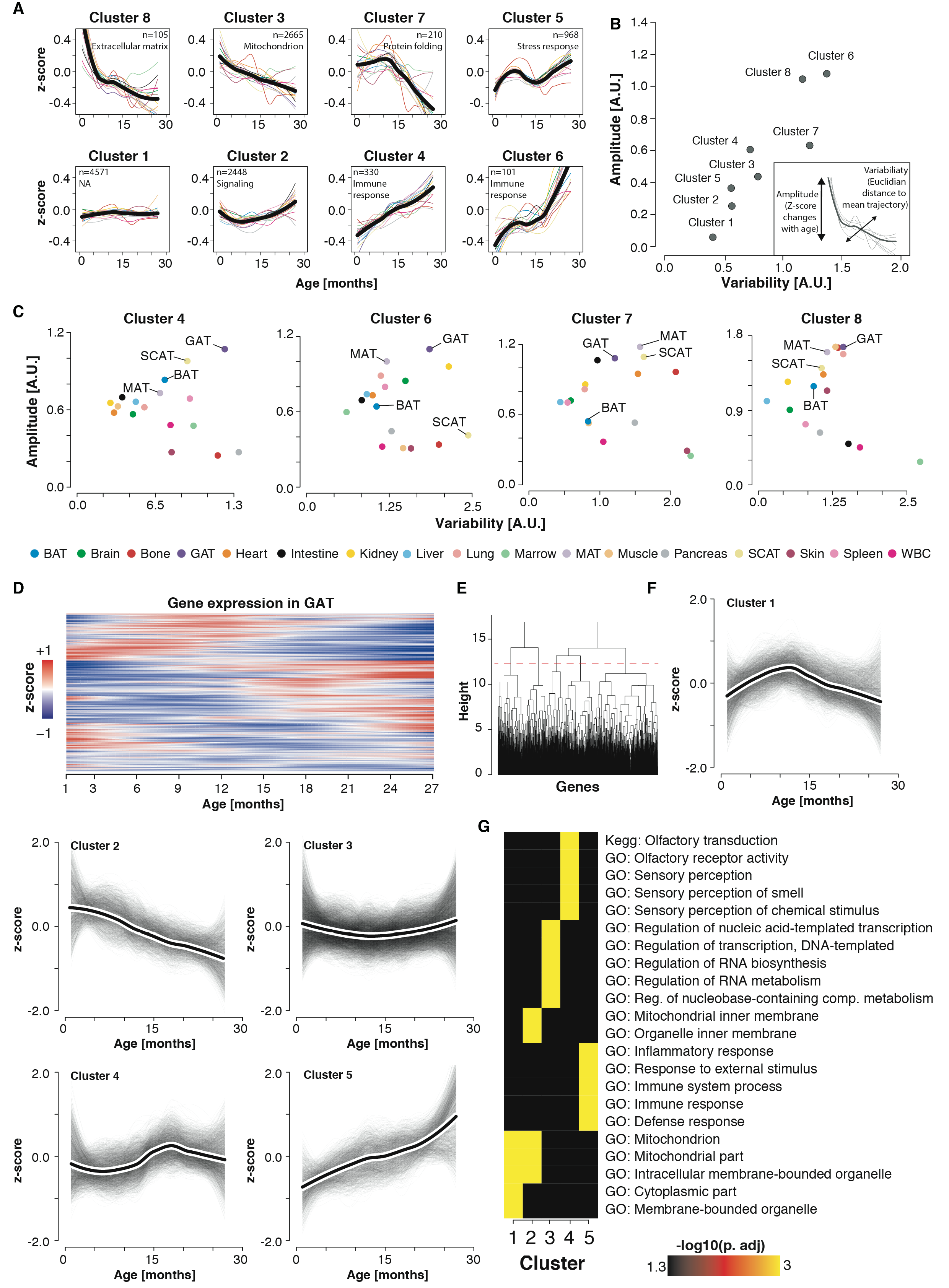
Aging gene expression dynamics across organs. (A) Whole-organism gene trajectories clustering. The average trajectory for each gene across all 17 organs was calculated. Whole-organism gene trajectories were grouped into 10 clusters, 8 clusters with more than 100 genes are represented. The number of genes and the top functionally enriched pathway for each cluster are reported. Cluster trajectories are indicated in black and average trajectories for each organ are represented by colors. (B) Identification of stable and variable clusters between organs. For each cluster in (A), an amplitude (absolute value of z-score change of the average trajectory between 1 and 27mos) and variability index (average Euclidian distance between the whole-organism trajectory and each organ-specific trajectory) were calculated. (C) Variability and amplitude index detailed per cluster. The 4 clusters changing the most in (B) are represented and adipose tissues are indicated since their changes are consistently among the top in the different clusters. (D) Focused clustering of GAT trajectories. Heatmap representing gene trajectories in GAT. Unsupervised hierarchical clustering was used to group genes with similar trajectories. (E) Clustering dendrogram and cut-off used to define 5 independent clusters in GAT. Aging trajectories for the 17 organs are available in Extended Data Figure 3. (F) Gene and cluster trajectories. Gene trajectories of the 5 clusters in (E) are represented in grey. Black lines surrounded by white represent the average trajectory for each cluster. (G) Enrichment for genes in functional categories. Pathway enrichment was tested using GOs, Reactome and KEGG databases for clusters in (E). The top 5 pathways for each cluster are shown. Pathways analysis for the 17 organs are available in Extended Data Figure 4.

Each cluster contains genes with similar global trajectories, but organ-specific differences in phase and magnitude suggest similar processes undergo unique dynamics. For each cluster we assigned an amplitude (absolute z-score change of the mean trajectory between 1mo and 30mo) and variability index (a measure of the spread from the mean trajectory) (Figure 2B). This revealed that clusters with the largest amplitudes also show the strongest organ-specific behavior, with adipose tissues prominently featured in clusters 4 and 6 (immune response), cluster 7 (protein folding), and cluster 8 (extracellular matrix) (Figure 2C). The life-long increase in immune response pathways is especially striking when organs are analyzed independently, especially for adipose tissues like GAT (Figure 2D-G, Extended Data Figure 4, Extended Data Figure 5, Extended Data Table 2).

These consistent expression shifts in whole organs may be driven by cell-intrinsic changes with age, or by changes in cell composition, such as immune cell accumulation. Given that visceral fat expansion predicts morbidity and mortality^9^, we aimed to discover the origin of the age-related adipose inflammatory signature by using single-cell RNA-sequencing (scRNA-seq) data from *Tabula Muris Senis*^10,11^. We identified increasing numbers of gonadal adipose tissue T and B cells with age, including a unique cluster of B cells present only in old mice (Figure 3A,B), consistent with increased expression of adaptive immune response genes in whole organs (Figure 2A,G). Unbiased screening of genes enriched in this population uncovered high Igj expression (Figure 3C, Extended Data Table 3). Considering we observed Igj differential expression in 11 of 17 whole organs (Figure 1G), we analyzed thousands of Cd79^+^ B cells across organs, revealing a unique cluster of Igj^high^ cells in both the FACS scRNA-seq and microfluidic droplet datasets, concordant with high expression of plasma B cell markers Xbp1 and Derl3 (Figure 3D, Extended Data Figure 6A,B,F,G). Relative to Igj^low^ cells, these cells show elevated unfolded protein response and endoplasmic reticulum stress pathways, characteristic of highly secretory plasma B cells (Figure 3E, Extended Data Figure 6C)^12^. Strikingly, these plasma B cells originate almost entirely from aged mice, and indeed accumulate across diverse organs (Figure 3F, Extended Data Figure 6D,E,H). Taking advantage of the high temporal resolution available with the whole-organ dataset, we also traced these cells across the lifespan via expression of Igj or Derl3. Indeed, we observed an initial increase of Igj in marrow, bone, and spleen, organs responsible for producing adaptive immune cells (Extended Data Figure 6I,J). Interestingly, Igj and Derl3 become subsequently elevated in kidney and GAT near 12 months of age, preceding Igj elevation in BAT, heart, and lung. The role of these cells or the specificity of the antibodies they produce is currently unknown, but it is tempting to speculate that they may contribute to the global increase in autoantibodies reported with aging^13^.

**Figure 3.**
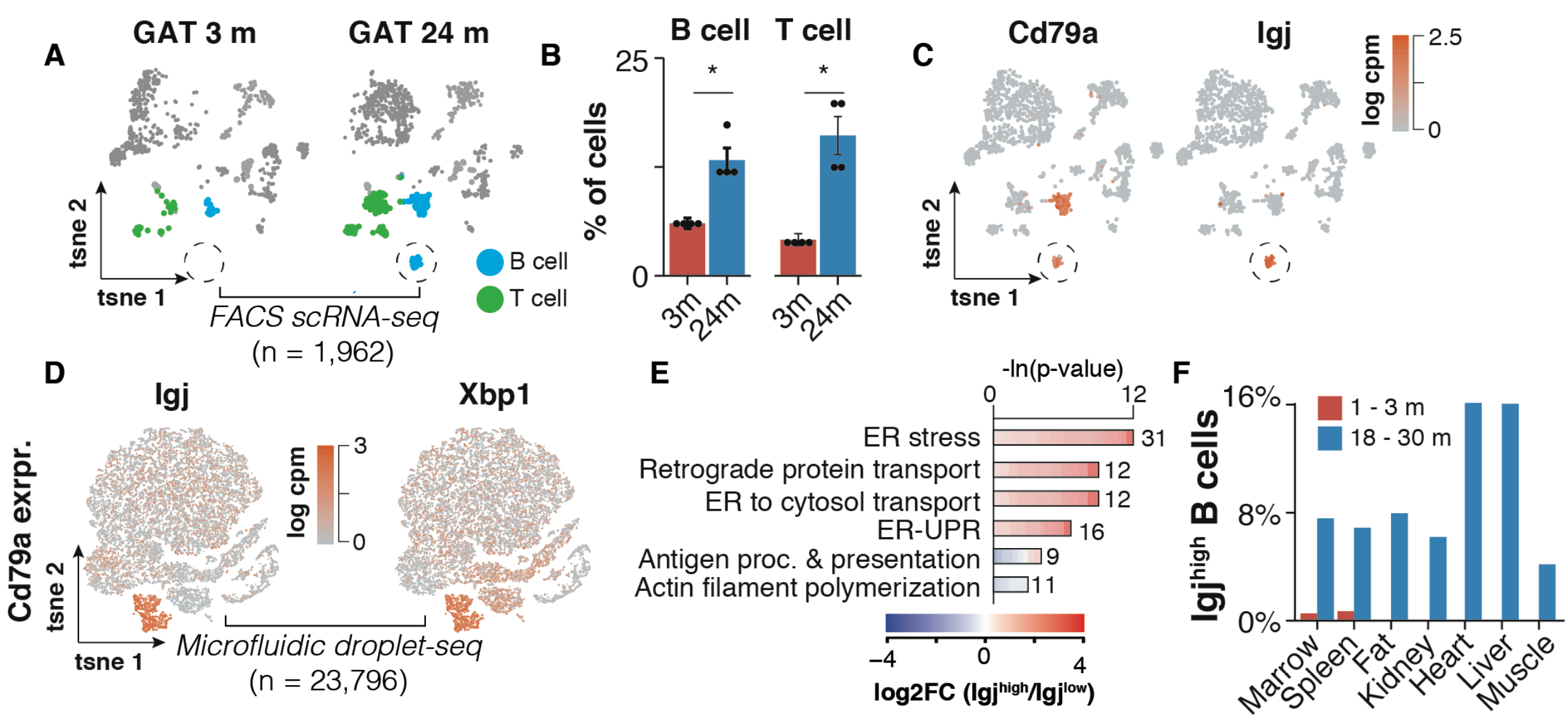
Single-cell transcriptomic analysis of Igjhigh plasma B cells. (A) t-SNE visualization of scRNA-seq data (FACS Smart-seq2) from the GAT stromal-vascular-fraction, split by age. B and T cells as annotated by the Tabula Muris Consortium are colored. A separate cluster of B cells present only in aged GAT is circled. (B) GAT B and T cells as a percentage of all analyzed cells. Means ± SEM, *** p<0.001, ** p<0.01, * p<0.05. (C) t-SNE as in (A) with young and old samples collapsed. Expression of B cell marker Cd79a and plasma B cell marker IgJ. (D) t-SNE visualization of scRNA-seq data (Droplet-seq) of all Cd79a-expressing cells present in the Tabula Muris Senis dataset (17 tissues), colored by the plasma B cell markers IgJ and Xbp1. (E) GO terms enriched among the top 300 marker genes of IgJ^high^ versus B cells. (F) Distribution of IgJ^high^ as percentages of Cd79a expressing cells per tissue.

Investigating organs individually can reveal detailed aging processes and even common phenotypes potentially susceptible to intervention. But aging occurs systemically, with the decline of one organ possibly inciting or accelerating dysfunction throughout the body. In part, this may be due to alterations to blood-borne factors which mediate intercellular and organ-organ communication. Spurred by heterochronic parabiosis experiments demonstrating rejuvenation^14,15^, we and others have identified plasma proteins with detrimental or rejuvenating functions in aging brain, muscle, pancreas, bone, and other organs^16^. However, the origins of these factors remain largely unknown.

Here, we attempted to uncover organs contributing to age-related changes in the plasma proteome by correlating plasma protein age trajectories with their corresponding gene expression trajectories in each organ (Figure 4A,B). This analysis reveals 25 plasma proteins correlated (Spearman correlation coefficient > 0.6) with gene expression in at least one organ, totaling 35 unique plasma protein/organ pairs. We discovered remarkably high correlation for several, such as vascular cell adhesion molecule-1 (Vcam1) in the kidney and fibroblast growth factor 10 (Fgf10) in the spleen, and other notable pairs such as glial fibrillary acidic protein (Gfap) and the brain. Especially interesting are Vcam1 and periostin (Postn), which both show exceptional correlation across several organs (Figure 4C-J). Vcam1 was recently implicated as a critical mediator of brain aging by old plasma^17^, and the loss of Postn in adipose tissues contributes to impaired lipid metabolism^18^. Throbospondin-4 (Thbs4), here decreasing with age and highly correlated with gene expression in muscle, was also recently discovered as a young blood-enriched protein that promotes synapse formation^19^. Interestingly, white adipose tissues emerge from this analysis as well, with 5 plasma proteins highly correlated with gene expression in visceral MAT and GAT, and 3 in SCAT (Figure 4A). Surprisingly, limb muscle which shows a modest number of DEGs displays 7 plasma proteins correlated with gene expression across the lifespan, including Postn, bone morphogenic protein-1 (Bmp1), matrix metalloprotease-2 (Mmp2), and other extracellular matrix (EM)-associated proteins (Extended Data Figure 7A). EM-associated proteins actually constitute of a majority of the 25 plasma proteins we identified (Extended Data Figure 7B). Future research will determine if these gene expression changes do indeed contribute to age-related differences in the plasma proteome, or if they mediate other processes like immune cell adhesion and infiltration.

**Figure 4.**
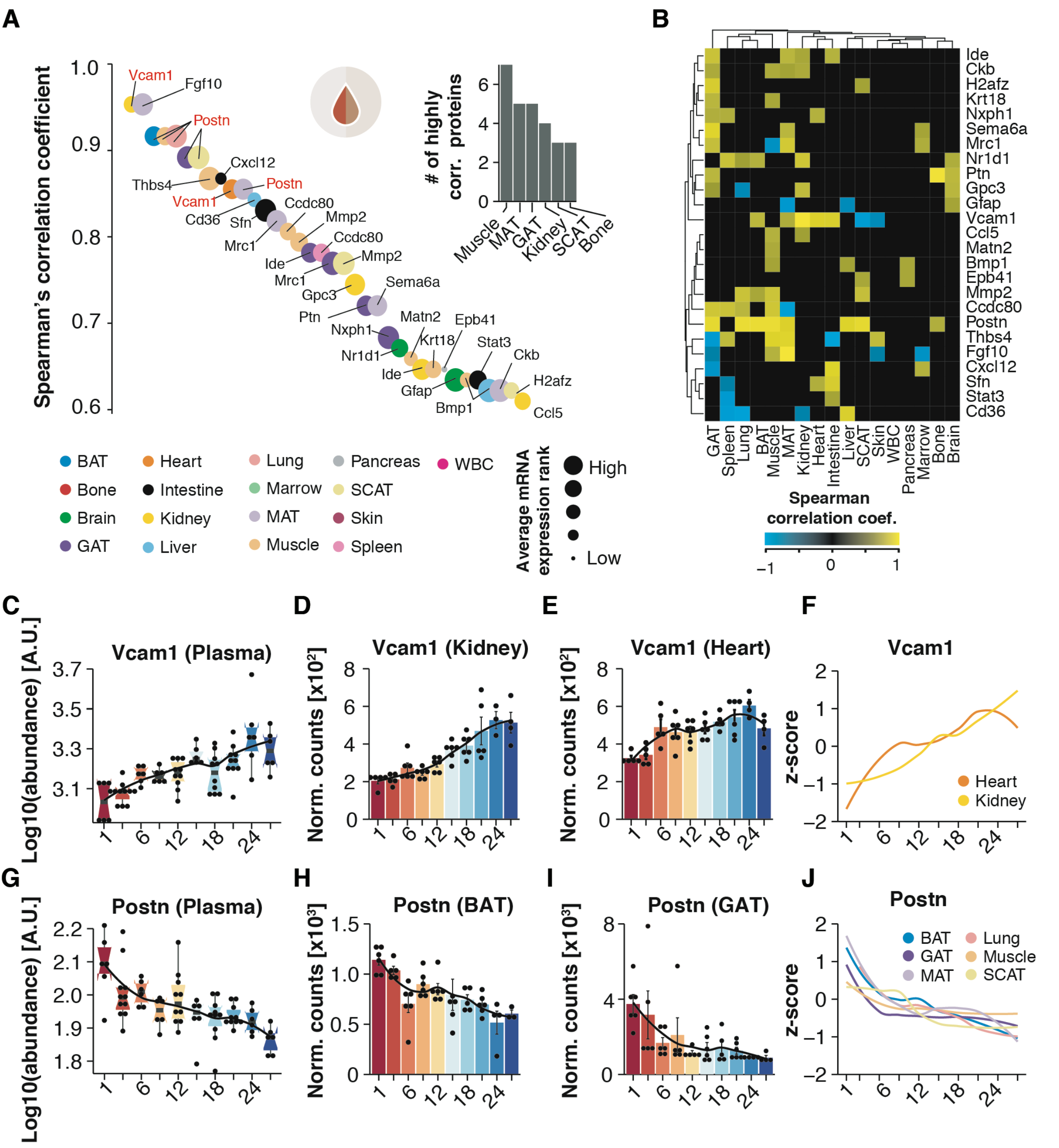
Plasma protein correlation with organ-specific gene expression. (A) Spearman correlation coefficient (≥ 0.6) between plasma proteins and corresponding organ-specific gene expression, colored by organ type. Dot size corresponds to average gene expression across tissues. Top right: bar graph of the number of proteins correlated with gene expression in the top 6 organs. (B) Heatmap showing correlation coefficients for the top 25 plasma proteins in (A) across all organs. (C) log-transformed plasma protein abundance of Vcam1. (D,E) Vcam1 mRNA expression (RNA-seq) in (D) Kidney and (E) Heart. LOESS regression is indicated by black line. (F) Z-transformed, smoothed gene expression trajectory of Vcam1, colored by tissue. (G) log-transformed plasma protein abundance of Postn. (H,I) Postn mRNA expression (RNA-seq) in (D) BAT and (E) GAT. LOESS regression is indicated by black line. (F) Z-transformed, smoothed gene expression trajectory of Postn, colored by tissue. Means ± SEM.

Altogether, our transcriptomic and plasma proteomic dataset offers unprecedented temporal resolution across the entire mouse lifespan for all major organs, and can serve as a fundamental resource to biologists across disciplines. With rejuvenation strategies like senescent cell ablation (senolytics), nutrient sensing manipulation (rapamycin and metformin), and plasma proteome alteration rapidly emerging, we need an improved understanding of where and when to apply these therapies^20^. This organism-wide characterization of aging dynamics may aid such therapeutic development, and the insights into circadian rhythm disruption, plasma cell accumulation, and adipose decline suggest avenues for renewed focus.

**Supplementary Information** is available in the online version of the paper.

## Acknowledgements

We thank members of the Wyss-Coray laboratory for volunteering help during organ collection and processing: Elizabeth Berber^3^, Kyle Brewer^2^, Betty Chang^3^, Julia Marschallinger^2^, Liana Bonanno^2^, Jerry Sun^2^, Maria F. Lugo-Fagundo^3^, Andrew Yang^2^, M. Windy McNerny^6,7^. We also thank the members of the Wyss-Coray laboratory and the Chan-Zuckerberg Biohub for feedback and support, and H. Zhang and K. Dickey for laboratory management. This work was funded by the Department of Veterans Affairs (BX004599 to T.W.-C.), the National Institute on Aging (R01-AG045034 and DP1-AG053015 to T.W.-C.), the NOMIS Foundation (T.W.-C.), The Glenn Foundation for Medical Research (T.W.-C.), and the Wu Tsai Neurosciences Institute (T.W.-C.).

## Author Contributions

N.S., B.L., S.R.Q., and T.W.-C. conceptualized the study. N.S., O.H., B.L., and T.W.-C. conceptualized the analysis. O.H. and B.L. with contributions from A.K., T.F., and R.P. conducted the analysis. N.S., S.E.L., D.P.L., M.E.Z., H.Z., and D.B., collected samples and extracted RNA. S.H., A.Z., and W.T. conducted cDNA and library preparation. P.M. created the Shiny web interface. R.S. and M.T. performed sequencing and library quality control. S.H., A.O.P., J.W., and A.M. processed raw sequencing data. The Tabula Muris Consortium generated the single-cell sequencing database. N.S., B.L., O.H., T.W.-C., and S.R.Q. wrote and edited the manuscript. T.W.-C., S.R.Q., S.D., N.F.N. and J.K. supervised the work.

## Author Information

Reprints and permissions information is available at www.nature.com/reprints. The authors declare no competing financial interests. Readers are welcome to comment on the online version of the paper. Correspondence to.

## Reviewer Information

*Nature* thanks the anonymous reviewers for their contributions to the peer review of this work.

## The Tabula Muris Consortium

**Overall Coordination:** Angela Oliveira Pisco^1^, Nicholas Schaum^2^, Aaron McGeever^1^, Jim Karkanias^1^, Norma F. Neff^1^, Spyros Darmanis^1^*, Tony Wyss-Coray^3-5^*, and Stephen R. Quake^1,6^*

**Organ collection and processing:** Jane Antony^2^, Ankit S. Baghel^2^, Isaac Bakerman^2,7,8^, Ishita Bansal^3^, Elizabeth Berber^5^, Daniela Berdnik^5^, Biter Bilen^3,4^, Liana Bonanno^3^, Kyle Brewer^3^, Douglas Brownfield^9^, Corey Cain^10^, Betty Chang^5^, Michelle B. Chen^4^, Stephanie D. Conley^2^, Spyros Darmanis^1^, Aaron Demers^2^, Kubilay Demir^2,11^, Antoine de Morree^3,4^, Tessa Divita^1^, Haley du Bois^5^, Laughing Bear Torrez Dulgeroff^2^, Hamid Ebadi^1^, F. Hernán Espinoza^9^, Matt Fish^2,11,12^, Qiang Gan^3,4^, Benson M. George^2^, Astrid Gillich^9^, Foad Green^1^, Geraldine Genetiano^1^, Xueying Gu^12^, Gunsagar S. Gulati^2^, Michael Seamus Haney^3^, Yan Hang^12^, Shayan Hosseinzadeh^1^, Albin Huang^4^, Tal Iram^4^, Taichi Isobe^1^, Feather Ives^2^, Robert Jones^3^, Kevin S. Kao^2^, Guruswamy Karnam^13^, Aaron M. Kershner^2^, Nathalie Khoury^3^, Bernhard M. Kiss^2,14^, William Kong^2^, Maya E. Kumar^15,16^, Jonathan Lam^12^, Davis P. Lee^6^, Song E. Lee^4^, Olivia Leventhal^5^, Guang Li^17^, Qingyun Li^18^, Ling Liu^3,4^, Annie Lo^1^, Wan-Jin Lu^1,9^, Maria F. Lugo-Fagundo^5^, Anoop Manjunath^1^, Julia Marschallinger^3^, Andrew P. May^1^, Ashley Maynard^1^, Marina McKay^1^, M. Windy McNerney^32,33^,Ross J. Metzger^19,20^, Marco Mignardi^1^, Dullei Min^21^, Ahmad N. Nabhan^9^, Norma F. Neff^1^, Katharine M. Ng^3^, Joseph Noh^2^, Rasika Patkar^13^, Weng Chuan Peng^12^, Lolita Penland^1^, Robert Puccinelli^1^, Eric J. Rulifson^12^, Nicholas Schaum^2^, Michael Seamus Haney^3^, Shaheen S. Sikandar^2^, Rahul Sinha^2,22-24^, Rene V. Sit^1^, Daniel Staehli^3^, Krzysztof Szade^2,25^, Weilun Tan^1^, Cristina Tato^1^, Krissie Tellez^12^, Kyle J. Travaglini^9^, Carolina Tropini^26^, Lucas Waldburger^1^, Linda J. van Weele^2^, Michael N. Wosczyna^3,4^, Jinyi Xiang^2^, Soso Xue^3^, Andrew C. Yang^3^, Lakshmi P. Yerra^5^, Justin Youngyunpipatkul^1^, Fabio Zanini^3^, Macy E. Zardeneta^6^, Fan Zhang^19^, Hui Zhang^5^, Lu Zhou^18^

**Library preparation and sequencing:** Spyros Darmanis^1^, Shayan Hosseinzadeh^1^, Ashley Maynard^1^, Norma F. Neff^1^, Lolita Penland^1^, Rene V. Sit^1^, Michelle Tan^1^, Weilun Tan^1^, Alexander Zee^1^

**Computational Data Analysis:** Oliver Hahn^3^, Lincoln Harris^1^, Benoit Lehallier^3^, Aaron McGeever^1^, Angela Oliveira Pisco^1^, Róbert Pálovics^30^

**Cell Type Annotation:** Jane Antony^2^, Biter Bilen^3,4^, Weng Chuan Peng^12^, Spyros Darmanis^1^, Antoine de Morree^3,4^, Oliver Hahn^3^, Yan Hang^12^, Shayan Hosseinzadeh^1^, Tal Iram^4^, Taichi Isobe^1^, Aaron M. Kershner^1^, Jonathan Lam^12^, Guang Li^17^, Qingyun Li^18^, Ling Liu^3,4^, Wan-Jin Lu^1,9^, Ashley Maynard^1^, Dullei Min^21^, He Mu^31^, Ahmad N. Nabhan^9^, Patricia K. Nguyen^2,7,8,17^, Angela Oliveira Pisco^1^, Nicholas Schaum^2^, Shaheen S. Sikandar^2^, Rahul Sinha^1,22-24^, Rene Sit^1^, Michelle Tan^1^, Weilun Tan^1^, Kyle J. Travaglini^9^, Margaret Tsui^13^, Bruce M. Wang^13^, Linda J. van Weele^2^, Michael N. Wosczyna^3,4^, Jinyi Xiang^2^, Alexander Zee^1^, Lu Zhou^18^

**Principal Investigators:** Ben A. Barres^18^, Philip A. Beachy^2,9,11,12^, Charles K. F. Chan^28^, Michael F. Clarke^2^, Spyros Darmanis^1^, Kerwyn Casey Huang^2,3,26^, Jim Karkanias^1^, Seung K. Kim^12,29^, Mark A. Krasnow^9,11^, Maya E. Kumar^15,16^, Christin S. Kuo^9,11,21^, Ross J. Metzger^19,20^, Norma F. Neff^2^, Roel Nusse^9,11,12^, Patricia K. Nguyen^2,7,8,17^, Thomas A. Rando^3-5^, Justin Sonnenburg^26^, Bruce M. Wang^13^, Kenneth Weinberg^21^, Irving L. Weissman^2,22-24^, Sean M. Wu^2,7,17^, Stephen R. Quake^2,6^, Tony Wyss-Coray^3-5^

^1^ Chan Zuckerberg Biohub, San Francisco, California, USA

^2^ Institute for Stem Cell Biology and Regenerative Medicine, Stanford University School of Medicine, Stanford, California, USA

^3^ Department of Neurology and Neurological Sciences, Stanford University School of Medicine, Stanford, California, USA

^4^ Paul F. Glenn Center for the Biology of Aging, Stanford University School of Medicine, Stanford, California, USA

^5^ Center for Tissue Regeneration, Repair, and Restoration, V.A. Palo Alto Healthcare System, Palo Alto, California, USA

^6^ Department of Bioengineering, Stanford University, Stanford, California, USA

^7^ Stanford Cardiovascular Institute, Stanford University School of Medicine, Stanford, California, USA

^8^ Department of Medicine, Division of Cardiology, Stanford University School of Medicine, Stanford, California, USA

^9^ Department of Biochemistry, Stanford University School of Medicine, Stanford, California, USA

^10^ Flow Cytometry Core, V.A. Palo Alto Healthcare System, Palo Alto, California, USA

^11^ Howard Hughes Medical Institute, USA

^12^ Department of Developmental Biology, Stanford University School of Medicine, Stanford, California, USA

^13^ Department of Medicine and Liver Center, University of California San Francisco, San Francisco, California, USA

^14^ Department of Urology, Stanford University School of Medicine, Stanford, California, USA

^15^ Sean N. Parker Center for Asthma and Allergy Research, Stanford University School of Medicine, Stanford, California, USA

^16^ Department of Medicine, Division of Pulmonary and Critical Care, Stanford University School of Medicine, Stanford, California, USA

^17^ Department of Medicine, Division of Cardiovascular Medicine, Stanford University, Stanford, California, USA

^18^ Department of Neurobiology, Stanford University School of Medicine, Stanford, CA USA

^19^ Vera Moulton Wall Center for Pulmonary and Vascular Disease, Stanford University School of Medicine, Stanford, California, USA

^20^ Department of Pediatrics, Division of Cardiology, Stanford University School of Medicine, Stanford, California, USA

^21^ Department of Pediatrics, Stanford University school of Medicine, Stanford, California, USA

^22^ Department of Pathology, Stanford University School of Medicine, Stanford, California, USA

^23^ Ludwig Center for Cancer Stem Cell Research and Medicine, Stanford University School of Medicine, Stanford, California, USA

^24^ Stanford Cancer Institute, Stanford University School of Medicine, Stanford, California, USA

^25^ Department of Medical Biotechnology, Faculty of Biophysics, Biochemistry and Biotechnology, Jagiellonian University, Poland

^26^ Department of Microbiology & Immunology, Stanford University School of Medicine, Stanford, California, USA

^27^ Department of Biochemistry and Biophysics, University of California San Francisco, San Francisco, California USA

^28^ Department of Surgery, Division of Plastic and Reconstructive Surgery, Stanford University, Stanford, California USA

^29^ Department of Medicine and Stanford Diabetes Research Center, Stanford University, Stanford,

California USA

^30^ Department of Computer Science, Stanford University, Stanford, California USA

^31^ Department of Physiology, University of California, San Francisco, CA 94158

^32^ Mental Illness Research Education and Clinical Center, V.A. Palo Alto Healthcare System, Palo Alto, California, USA

^33^ Department of Psychiatry, Stanford University School of Medicine, Stanford, California, USA

**Extended Data Figure 1.**
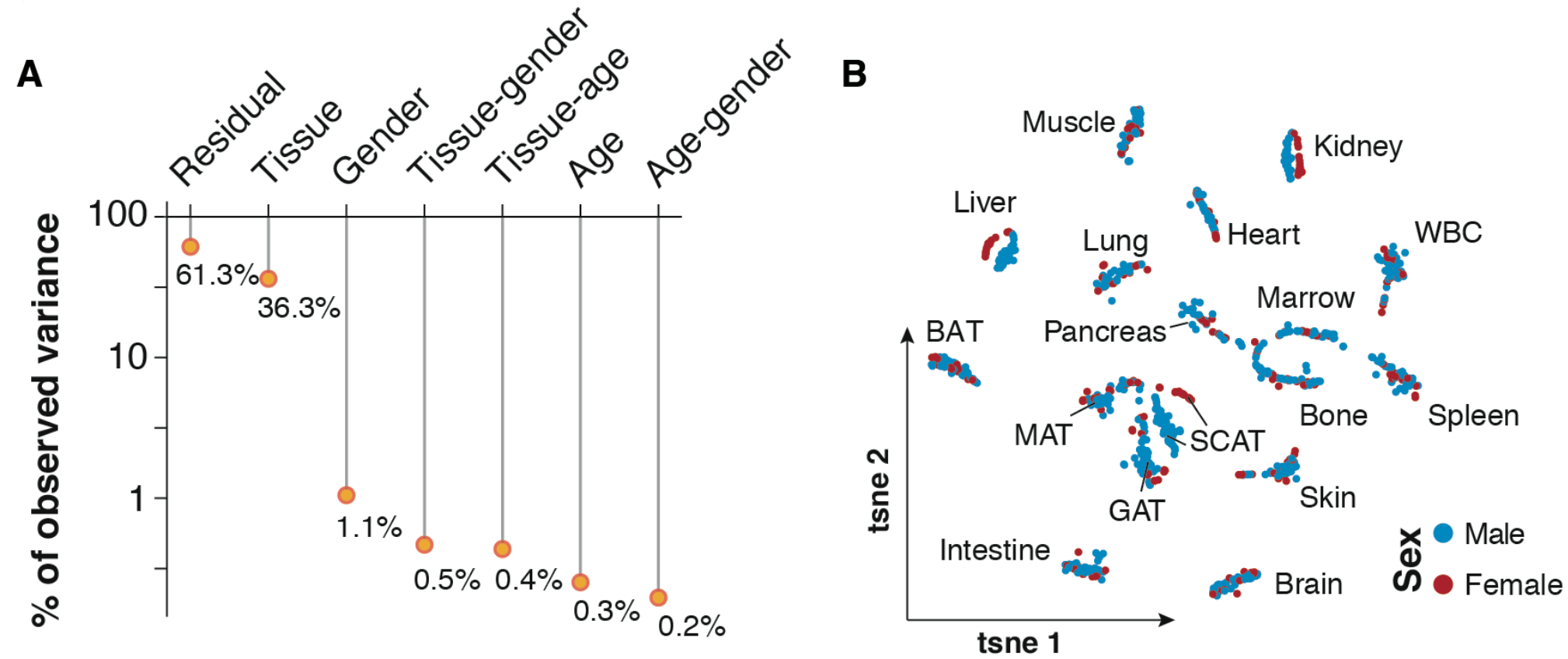
Gene expression variance analysis. (A) Visualization of the Principle Variance Component Analysis, displaying the gene expression variance explained by residuals (i.e. biological and technical noise) or experimental factor such as tissue, age, gender and respective combinations. (B) t-SNE visualization of all samples, colored by sex.

**Extended Data Figure 2.**
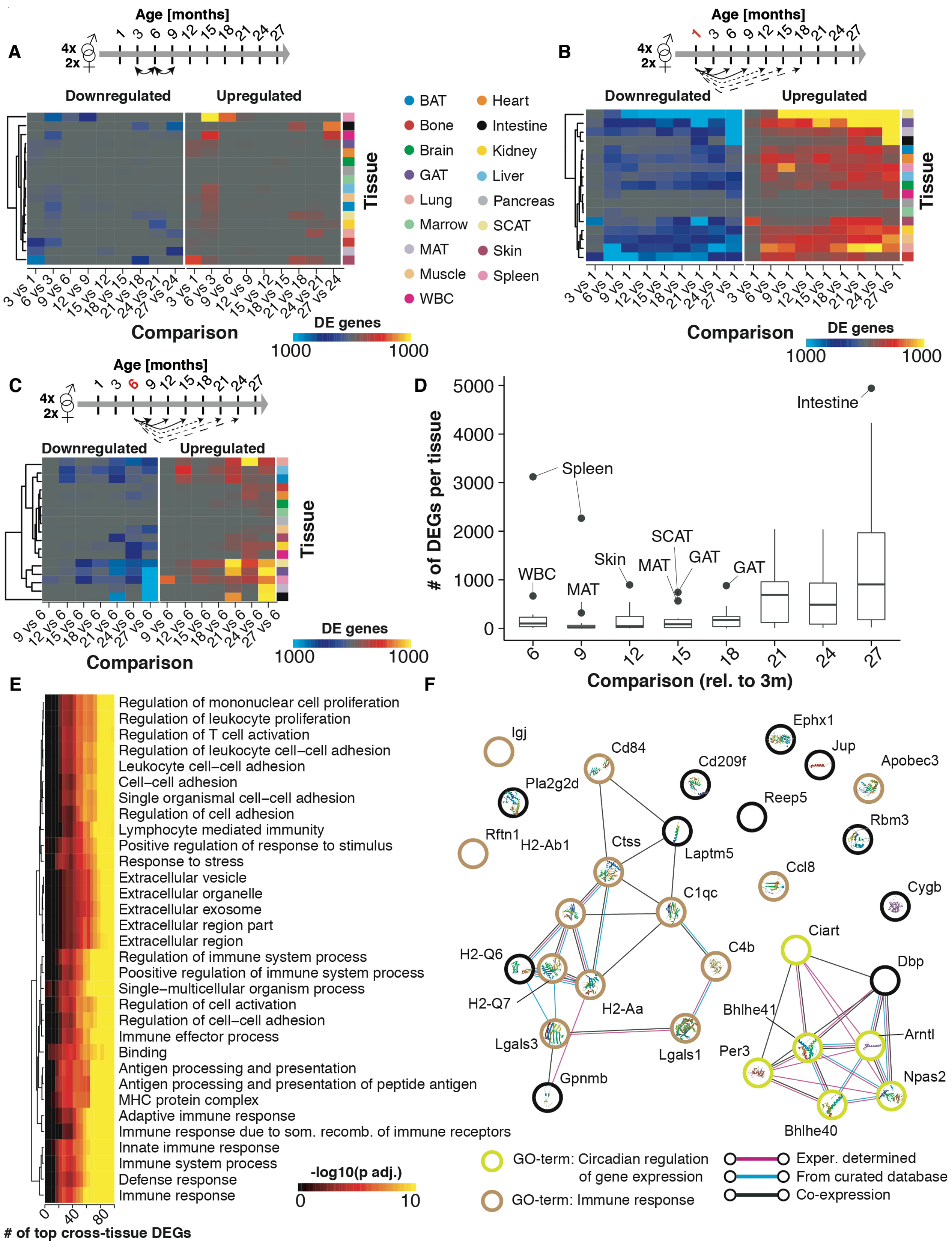
Validation of differential gene expression analysis. (A) Heatmap representation displaying the number of differentially expressed genes per tissue when conducting pairwise analysis consecutive sampling timepoints. (B) Heatmap representation displaying the number of differentially expressed genes per tissue for pairwise comparisons with a 1mo reference. (C) Heatmap representation displaying the number of differentially expressed genes per tissue for pairwise comparisons with a 6mo reference. (D) Boxplot representation displaying the number of differentially expressed genes per tissue for pairwise comparisons with a 3mo reference. Outliers allow detection of tissues undergoing exceptionally strong expression shifts at a given age. (E) Enrichment for genes in functional categories. Genes were ranked by number of differentially expressed genes per tissue for pairwise comparisons with a 3mo reference. Pathway enrichment was tested using GOs, Reactome and KEGG databases (F) STRING analysis of the top 30 genes in Figure 1G.

**Extended Data Figure 3.**
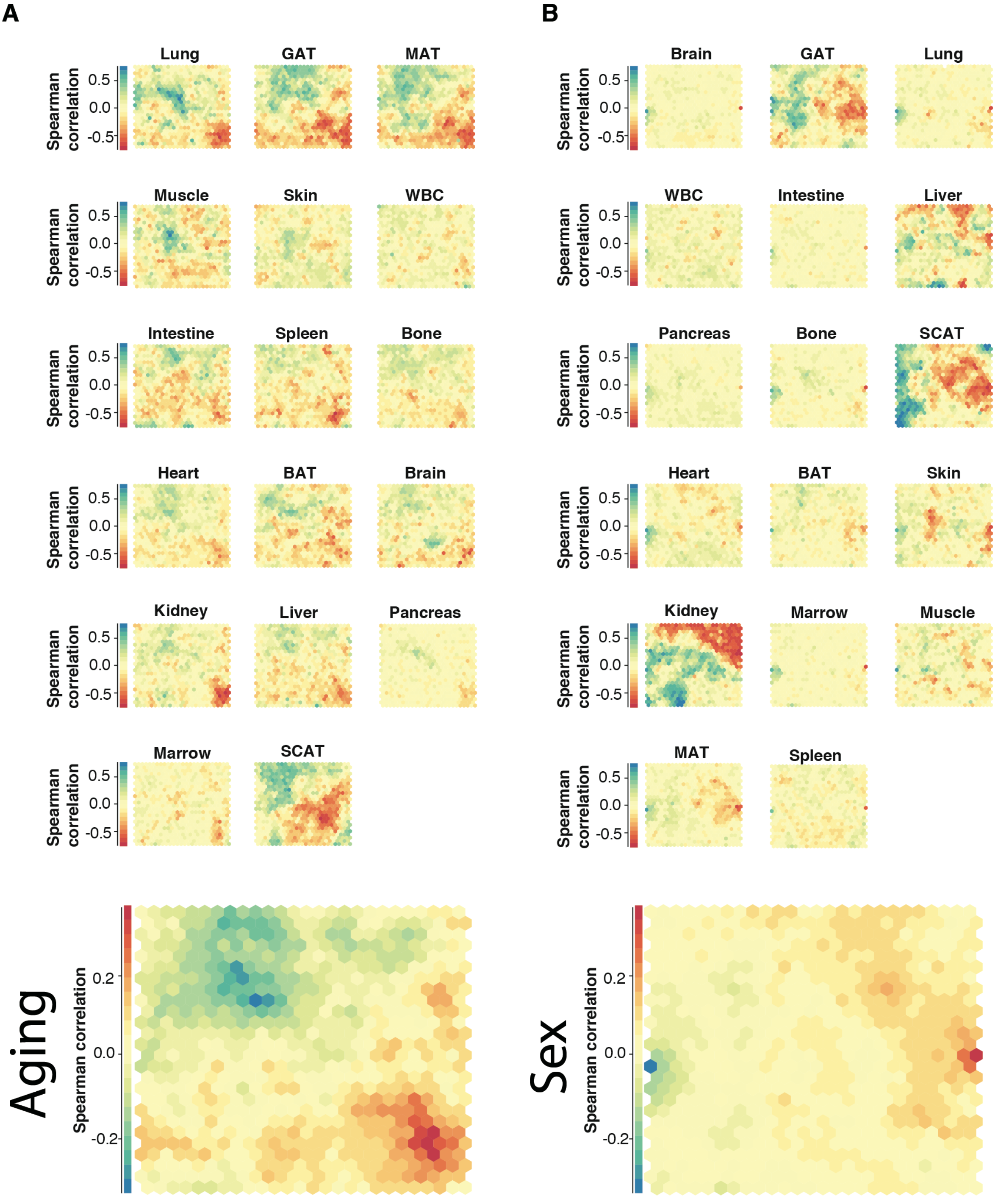
Self-organizing maps of gene correlation with age and sex. Self-organizing maps (SOMs) were generated from transcriptome-wide gene expression correlation (Spearman’s rank correlation coefficient) of each gene with age (A) and sex (B). Genes with similar correlation are mapped to the same cell, and cells grouped by similarity. The SOM cell layout is common across organs, with the average across all organs at bottom.

**Extended Data Figure 4.**
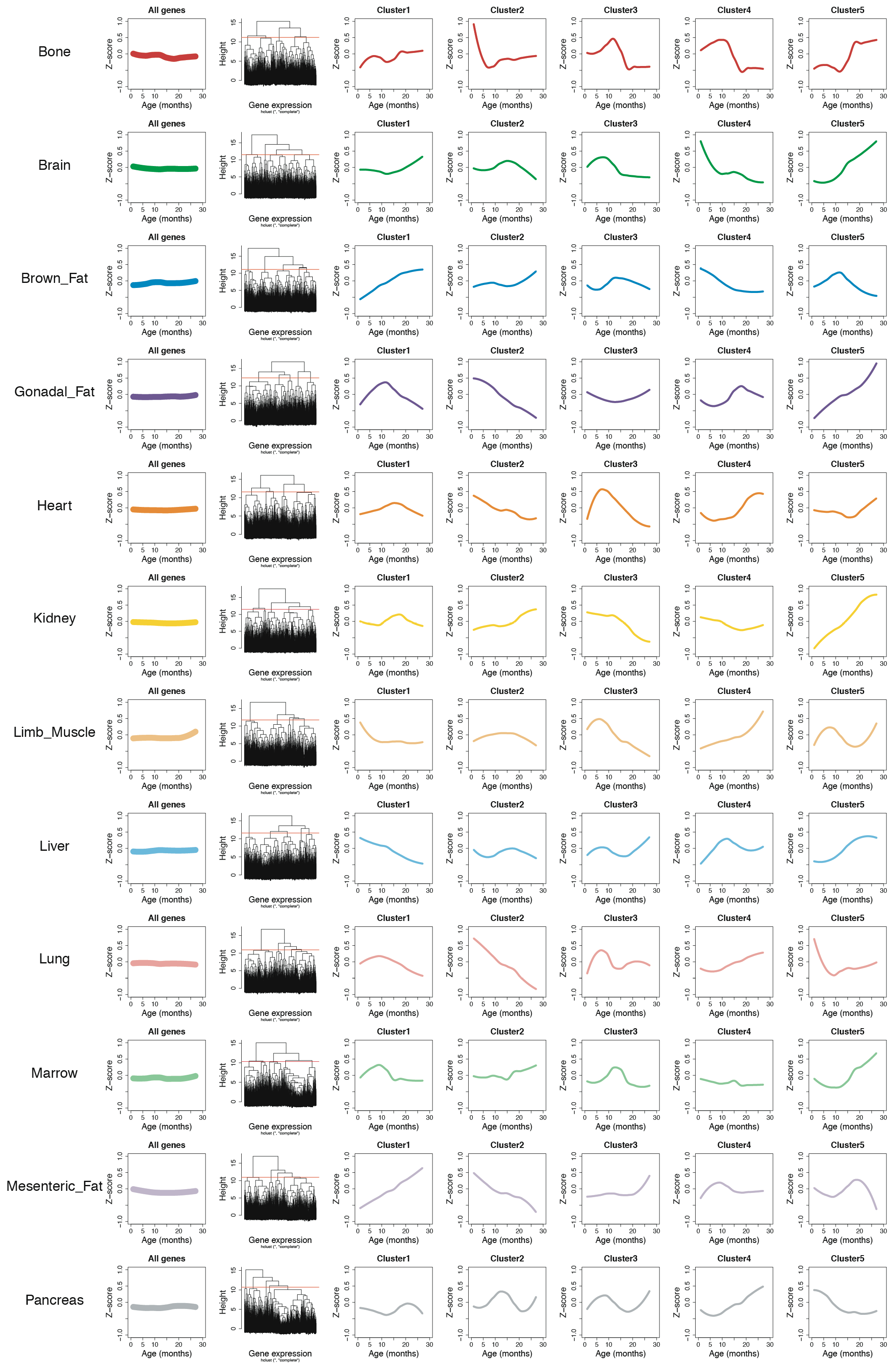
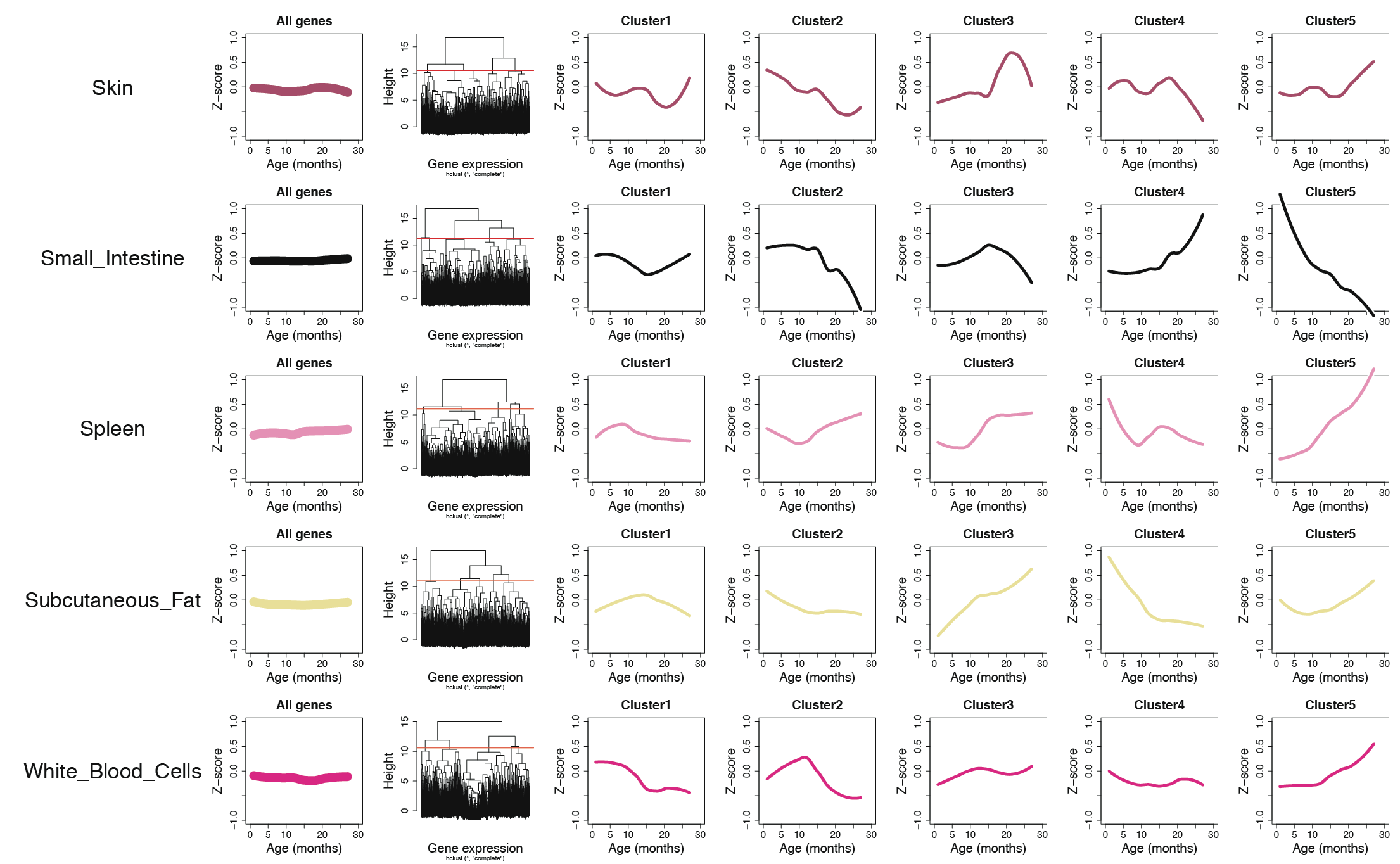
Organs-specific gene expression dynamics. For each of the 17 organs (rows), the average trajectory is represented in the 1^st^ column and unsupervised hierarchical clustering was used to group genes with similar trajectories (columns 2). Five cluster were used (columns 3-7) for further analysis.

**Extended Data Figure 5.**
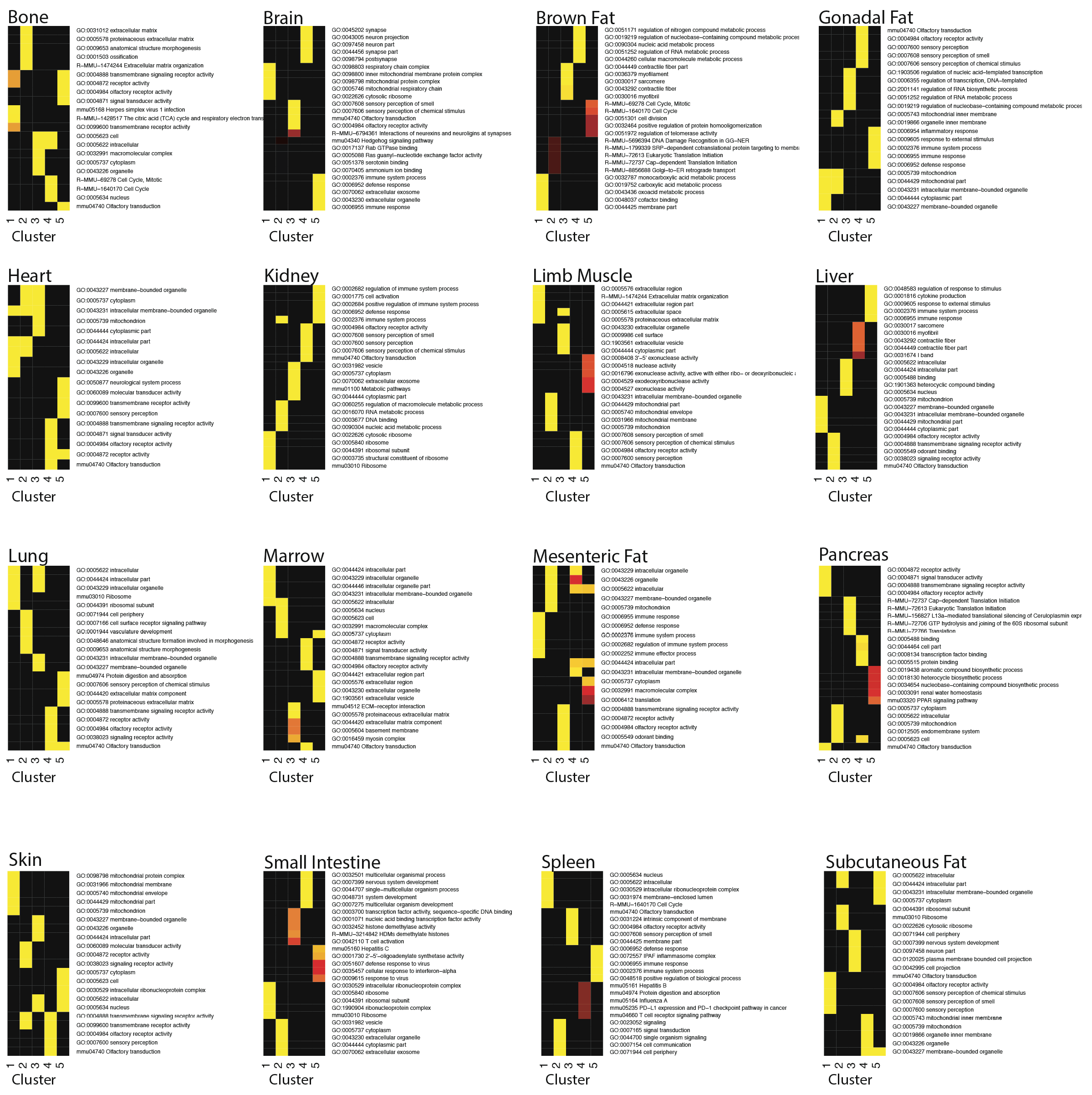
Pathways enrichment analysis of organs-specific clusters. Clusters in (Extended Data Figure 4) show enrichment for genes in functional categories. Pathway enrichment was tested using GOs, Reactome and KEGG databases. The top 5 pathways for each cluster are shown.

**Extended Data Figure 6.**
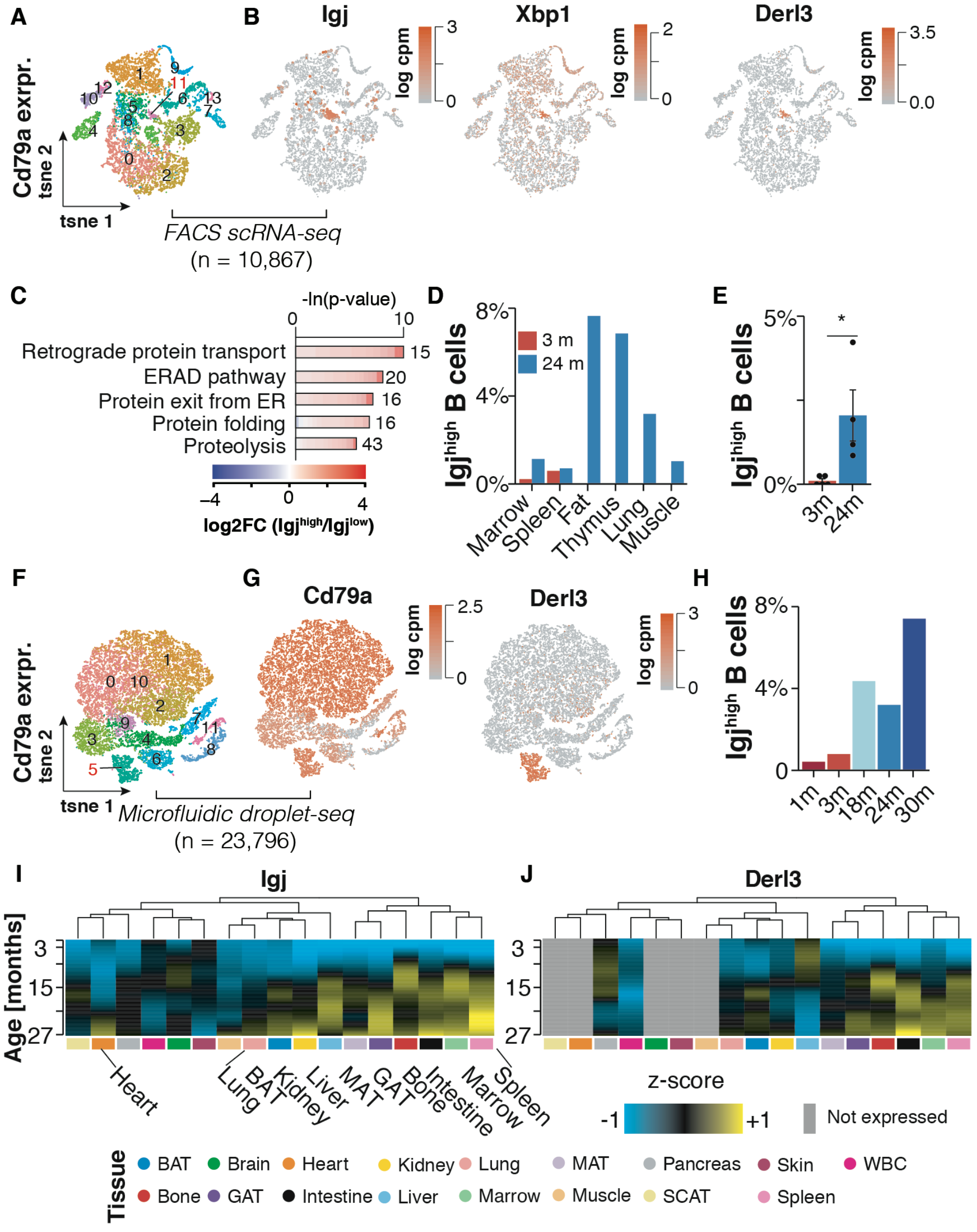
Identifying IgJhigh B cells with Smart-seq2 and Droplet-seq. (A) t-SNE visualization of scRNA-seq data (FACS Smart-seq2) of all Cd79a-expressing cells present in the Tabula Muris Senis dataset (17 tissues). Colored are clusters of cells as identified with the Seurat software toolkit. Highlighted is cluster 11, comprised of IgJ^high^ B cells. (B) t-SNE in (A) colored by the IgJ^high^ B cell markers IgJ, Xbp1 and Derl3. (C) GO terms enriched among the top 300 marker genes of IgJ^high^ versus B cells (FACS Smart-seq2). (D) Distribution of IgJ^high^ as percentages of Cd79a expressing cells per tissue. (E) Percentage of IgJ^high^ B cells of all Cd79a expressing cells across all tissues. Means ± SEM, *** p<0.001, ** p<0.01, * p<0.05. (F) t-SNE visualization of scRNA-seq data (Droplet-seq) of all Cd79a-expressing cells present in the Tabula Muris Senis dataset (17 tissues). Colored are clusters of cells as identified with the Seurat software toolkit. Highlighted is cluster 5, comprised of IgJ^high^ B cells. (G) t-SNE in colored by the B cell marker Cd79a and IgJ^high^ B cell marker Derl3. (H) Percentage of IgJ^high^ B cells of all Cd79a expressing cells across all tissues. (I, J) Heatmap visualization of the z-transformed expression trajectories of (I) IgJ and (J) per tissue as measured by mRNA-seq.

**Extended Data Figure 7.**
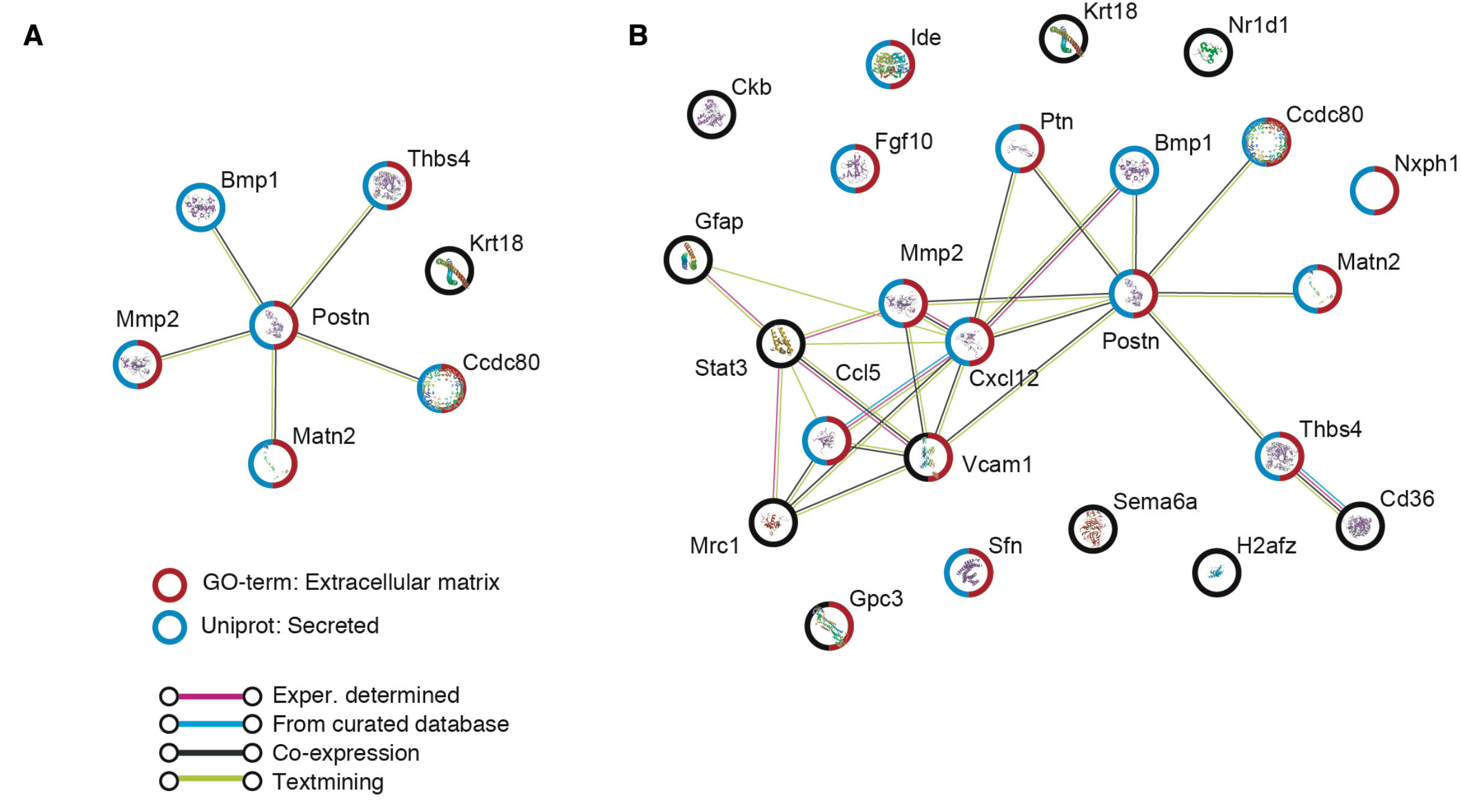
STRING analysis of top correlating plasma proteins. The top 7 plasma proteins correlated with gene expression in muscle (A), and the top 25 plasma proteins correlated with gene expression in any organs (B), colored by pathway.

## Methods

### Mice and Organ Collection

Male and virgin female C57BL/6JN mice were shipped from the National Institute on Aging colony at Charles River (housed at 67–73 °F) to the Veterinary Medical Unit (VMU; housed at 68–76 °F)) at the VA Palo Alto (VA). At both locations, mice were housed on a 12-h light/dark cycle, and provided food and water ad libitum. The diet at Charles River was NIH-31, and Teklad 2918 at the VA VMU. Littermates were not recorded or tracked, and mice were housed at the VA VMU for no longer than 2 weeks before euthanasia, with the exception of mice older than 18mos, which were housed at the VA VMU beginning at 18mos of age. After anaesthetization with 2.5% v/v Avertin, mice were weighed, shaved, and blood was drawn via cardiac puncture before transcardial perfusion with 20 ml PBS. Whole organs were then dissected in the following order: pancreas, spleen, brain, heart, lung, kidney, mesenteric adipose tissue, intestine (duodenum), gonadal adipose tissue, muscle (tibialis anterior), skin (dorsal), subcutaneous adipose tissue (inguinal pad), brown adipose tissue (interscapular pad), bone and bone marrow (femurs and tibiae). Mice were randomized and organs collected from 8:30am – 4pm over several days. Organs were immediately snap frozen on dry ice. All animal care and procedures were carried out in accordance with institutional guidelines approved by the VA Palo Alto Committee on Animal Research.

### Sample size, randomization, and blinding

No sample size choice was performed before the study. Randomization was performed in the case of mouse dissection order and during the preparation of 96-well plates for cDNA creation. Blinding was not performed: the authors were aware of all data and metadata-related variables during the entire course of the study.

### RNA isolation and preparation

Snap-frozen bone and skin was crushed on liquid nitrogen with a mortar and pestle. Snap-frozen whole organs, or crushed bone or skin, were placed in TRIzol and immediately homogenized with a TissueRuptor in 50ml conicals (see table for organ-specific details). Debris from the homogenate was pelleted in 1.5ml tubes at 12,000 x g for 5 minutes at 4°C. Supernatent was then transferred to a new 1.5ml tube where chloroform was added. After vortexing on max speed for 10 seconds, samples were transferred to 1.5ml or 2ml Phase Lock Gel tubes, where water was added before spinning at 12,000 x g for 5 minutes at 4°C. The aqueous phase was then transferred to a new tube, and after adding isopropanol, mixtures were vortexed at max speed for 10 seconds. Solutions were then run through RNeasy columns according to the manufacturer’s instructions, and eluted with the indicated volume of water. RNA was then quantified with a nanodrop, and frozen at −80°C.

| **Organ** | **RNeasy kit** | **TRIzol (ml)** | **Homogenate (ul)** | **CHCl<sub>3</sub> (ul)** | **H<sub>2</sub>O (ul)** | **Isopropanol (ul)** | **Elution (ul)** |
| --- | --- | --- | --- | --- | --- | --- | --- |
| **BAT** | mini | 1/50mg | 350 | 117 | 117 | 175 | 20 |
| **bone** | micro | 1.5 | 625 | 208 | 208 | 313 | 10 |
| **brain** | mini | 1/50mg | 625 | 208 | 208 | 313 | 30 |
| **GAT** | micro | 1/50mg | 1000 | 333 | 333 | 500 | 10 |
| **heart** | mini | 1/50mg | 625 | 208 | 208 | 313 | 30 |
| **kidney** | mini | 1/50mg | 300 | 100 | 100 | 150 | 30 |
| **muscle** | mini | 1/50mg | 400 | 133 | 133 | 200 | 20 |
| **liver** | mini | 1/50mg | 300 | 100 | 100 | 150 | 60 |
| **lung** | mini | 1/50mg | 625 | 208 | 208 | 313 | 30 |
| **marrow** | mini | 1/50mg | 900 | 300 | 300 | 450 | 20 |
| **MAT** | micro | 1/50mg | 1000 | 333 | 333 | 500 | 10 |
| **pancreas** | mini | 1/25mg | 200 | 67 | 67 | 100 | 30 |
| **SCAT (2X)** | micro | 1/50mg | 1000 | 333 | 333 | 500 | 10 |
| **skin** | mini | 1/50mg | 700 | 233 | 233 | 350 | 20 |
| **intestine** | mini | 1/50mg | 200 | 67 | 67 | 100 | 30 |
| **spleen** | mini | 1/50mg | 200 | 67 | 67 | 100 | 30 |
| **WBC** | mini | 3X vol | 625 | 208 | 208 | 313 | 20 |

### cDNA synthesis, library preparation, sequencing, and data processing

Methods including cDNA synthesis using the Smart-seq2 protocol^21^, library preparation using an in-house version of Tn5^22,23^, library pooling, quality control, sequencing, and data processing are provided at dx.doi.org/10.17504/protocols.io.2uvgew6

### Differential expression analysis

Samples with less than 4 mio uniquely mapped reads were censored to exclude low-coverage samples. Data visualization and analysis were performed using custom Rstudio scripts and the following Bioconductor packages: Rtsne, Deseq2^24^, topGO, and org.Mm.eg.db. In order to detect whether samples in a given tissue would show profound clustering by age, we calculated t-SNE maps based on the first principal component derived of the initial PCA step, i.e. the PC explaining most of the variance. To analyze significant expression changes with age, size factors and dispersion estimates were calculated for each tissue separately. We conducted differential expression analysis comparing samples from 3 months old mice to each consecutive time point, using age and sex as covariates. P-values were adjusted for multiple testing and genes with an adjusted p-value of below 0.05 were determined as statistically significant. In addition, we ran similar analyses using 1 month or 6 months old mice as reference.

To rank genes based on their regulation across tissues, we summarized in how many tissues a given gene would be called as significantly regulated in at least one comparison between samples from 3 months old mice and any following sampling time point.

### Gene expression trajectory analysis

To estimate genes trajectories during aging, normalized counts from DEseq2 were z-scored and LOESS (Locally weighted scatterplot smoothing) regression was fitted for each gene using the median expression per age group in each tissue. Whole-organism trajectory per gene was estimated using the average trajectory across the 17 tissues. Organism-wide analysis focused on 11403 genes expressed in all tissues (i.e. genes among the 15k most expressed genes in each tissue).

The distance matrix between whole-organism gene trajectories was computed using the euclidian distance and hierarchical clustering was performed using the complete method. We identified 10 clusters of genes changing with age, ranging from 1 to 4571 genes. Clusters 9 and 10 were excluded from further analysis as they included less than 10 genes.

To identify clusters changing the most between tissues, we computed an amplitude and variability index. The amplitude index corresponds to the z-score change (absolute value) of the average trajectory between 1 and 27mos. The variability index, which measures the spread of organ-trajectories, corresponds to the average Euclidian distance between each organ-specific trajectory and the organism-wide trajectory.

Reactome, KEGG and GO databases were queried to understand the biological functions of each cluster. We used the R TopGO^25^ package for GO analysis and the R clusterprofiler^26^ package for KEGG and Reactome analyses. The 11403 genes expressed in all tissues served as the background set of genes against which to test for over-representation. Since clusterprofiler requires EntrezID as input, we mapped Gene Symbols to EntrezID using the org.Mm.eg.db^27^ package. When Gene Symbols were mapped to multiple EntrezID, only the 1st EntrezID was used. Q-values were estimated using Benjamini–Hochberg approach for the different databases taken separately. In addition, for GO analysis, q-values were calculated for the three GOs classes (molecular function, cellular component, biological process) independently.

Organ-specific clustering were performed using the 15k most expressed genes per tissue. For each tissue, five clusters were considered for further analysis and the pathways analysis used the corresponding background set of genes against which to test for over-representation.

### Single-cell RNA-sequencing analysis (FACS scRNA-seq)

Pre-processed and annotated scRNA-seq data (FACS followed by Smart-seq2 protocol) from gonadal adipose tissue (3 and 24 months old) were obtained from the Tabula Muris Senis consortium. Given the lack of data from aged female mice, we focused our analyses on samples derived from male mice. Additionally, cells with less than 200 or more than 6,500 genes were excluded. This yielded 1,962 high quality cell transcriptomes derived from four young and four old biological replicates. Data visualization and analysis were performed using custom Rstudio scripts and the following Bioconductor packages: Seurat (version 3)^28^ and topGO. Data normalization and scaling was performed using Seurat’s built-in SCTransform^29^ function with default parameters. A shared-nearest-neighbors graph was constructed using the first 30 PC dimension before clustering cells using Seurat’s built-in FindClusters function with a resolution of 0.8 and default parameters. Annotations for B and T cells were adopted from the Tabula Muris Senis consortium. Cell numbers were normalized to the total number of detected cells and compared using standard t-tests. Igj^high^ B cells formed a separate cluster and were identified using Seurat’s FindMarkers function (parameters: only.pos=T min.pct=0.15 thresh.use=0.25, test=’MAST’).

In order to profile Igj^high^ B cells organism-wide, we obtained the complete and pre-processed scRNA-seq dataset (FACS followed by Smart-seq2 protocol) from the Tabula Muris Senis consortium – encompassing cells isolated from all major tissues. Focusing on data from male samples only, we filtered for cells showing detectable expression of the Cd79a gene (alpha chain of the B cell receptor). Cell transcriptomes were thus derived from four young and four old biological replicates. The resulting 10,867 cells were analyzed using Seurat, as described above. A shared-nearest-neighbors graph was constructed using the first 10 PC dimension before clustering cells using Seurat’s built-in FindClusters function with a resolution of 0.4 and default parameters. Igj^high^ B cells formed a separate cluster (cluster 11; 129 cells). To characterize Igj^high^-specific expression profile, Seurat’s FindMarkers function (parameters: only.pos=F min.pct=0.15 thresh.use=0.25, test=’MAST’) was run comparing the cluster of Igj^high^ B cells against all other Cd79a cells in the dataset. Functional enrichment analysis of the top 300 differentially expressed genes (sorted by adjusted p-value) were compared to all 1051 genes passing the filtering parameters for the test. Top-ranked GO terms were selected and visualized using the CellPlot package (https://github.com/dieterich-lab/CellPlot). The full-length GO terms were shortened to fit into the figure format; the complete table of significantly enriched GO terms and associated genes can be found in Extended Data Table 3.

### Single-cell RNA-sequencing analysis (Microfluidic droplet-seq)

The Tabula Muris Senis consortium encompasses scRNA-seq data generated with microfluidic droplets, allowing to profile more cells without prior selection of surface markers^30^. In order to profile Igj^high^ B cells organism-wide, we obtained the complete and pre-processed Droplet-seq dataset. Focusing on data from male samples only, we filtered for cells showing detectable expression of the Cd79a gene. Cell transcriptomes were derived from the following sampling timepoints: 2x 1 month old, 2x 3 months old, 2x 18 months, 4x 24 months and 3x 30 months. The resulting 23,796 cells were analyzed using Seurat, as described above. A shared-nearest-neighbors graph was constructed using the first 10 PC dimension before clustering cells using Seurat’s built-in FindClusters function with a resolution of 0.4 and default parameters. Igj^high^ B cells formed a separate cluster (cluster 5; 1,198 cells). To characterize Igj^high^-specific expression profile, Seurat’s FindMarkers function (parameters: only.pos=F min.pct=0.15 thresh.use=0.25, test=‘MAST’) was run comparing the cluster of Igj^high^ B cells against all other Cd79a cells in the dataset. Functional enrichment analysis of the top 300 differentially expressed genes (sorted by adjusted p-value) were compared to all 1886 genes passing the filtering parameters for the test. Top-ranked GO terms were selected and visualized using the CellPlot package (https://github.com/dieterich-lab/CellPlot). The full-length terms were shortened to fit into the figure format; the complete table of significantly enriched GO terms and associated genes can be found in Extended Data Table 3.

### Plasma proteomic analysis

All animal care and procedures were carried out in accordance with institutional guidelines approved by the VA Palo Alto Committee on Animal Research. Sixty-five male (n=5-6 per age group, 1mo/3mo/6mo/9mo/12mo/15mo/18mo/21mo/24mo/27mo/30mo) and 16 virgin female (n=4 per age group, 3mo/12mo/18mo/21mo) C57BL/6JN mice were shipped from the National Institute on Aging colony at Charles River (housed at 67–73 °F) to the Veterinary Medical Unit (VMU; housed at 68– 76 °F)) at the VA Palo Alto (VA). Mice were provided food (NIH-31 at Charles River, and Teklad 2918 at the VA VMU) and water at ad libitum. Mice were housed on a 12-h light/dark cycle at both places. Mice older than 18- months were housed at the VA VMU until they reached the experimental age. Mice younger than 18- months were housed for less than 2 weeks at the VA VMU. After anaesthetization with 2.5% v/v Avertin, blood was drawn via cardiac puncture. EDTA-plasma was isolated by centrifugation at 1,000g for 10 min at 4 °C. Samples were aliquoted, stored at −80 °C and sent on dry ice to SomaLogic Inc. (Boulder, Colorado, US).

The SomLogic platform is primarily designed to detect and measure human proteins. In order to reduce the influence of cross-species effects on our analysis, we first determined proteins in our dataset with high evolutionary conservation between mouse and humans. To this end, we downloaded the plain text file containing all homologies between mouse and human along with sequence identifiers for each species (HOM_MouseHumanSequence.rpt) from MGI (http://www.informatics.jax.org/). Next, reference protein sequences for human and mouse were downloaded from UniProt (https://www.uniprot.org/). Using the R “Biostrings” library a global pairwise sequence alignment has been carried out between the human and mouse sequences. Further, only sequences with identity of 80% across the whole alignment were included in the downstream analyses.

To determine the effect of age on the plasma proteome, Relative fluorescent units (RFUs) provided by Somalogic were log10- and linear models adjusted for age and sex were used. Type II sum of squares (SS) were calculated using R car package^31^ and q-values were estimated using Benjamini– Hochberg approach^32^.

Raw protein abundance data as measured by the Somalogic platform were scaled (z-transformed) before calculating the median across all replicates for each sampled age time point to obtain an average trajectory. Normalized RNA-seq counts after pre-processing with Deseq2 were transformed for each tissue alike. In order to compare plasma protein changes with shifts in mRNA expression in any tissue, we calculated pairwise Spearman’s rank correlation coefficients between a given protein trajectory with the expression trajectory for its’ corresponding gene in each tissue separately. Thus, a given protein could be correlated with expression changes in multiple tissues. In order to limit our analysis to mRNA /protein pairs reflecting robust changes, we filtered the resulting mRNA/protein correlations as follows: (1) The protein had to exhibit a sequence homology between human and mouse of at least 75%. (2) The protein had to change significantly with age according to the linear modeling analysis. (3) The corresponding gene had to be differentially expressed in the given tissue in at least one pairwise comparison between 3 months old mice and any consecutive time point. (4) The mRNA /protein profiles had to exhibit a Spearman’s rank correlation of at least 0.6. Given that genes can be expressed at differing levels across tissues, we additionally calculated average mRNA expression ranks for each gene. To this end, we ranked for a given gene each tissue on its’ average mRNA expression, based on Deseq2’s baseMean.

To investigate connectivity networks between top proteins correlated with organ gene expressed, we used String version 11.0, available at https://string-db.org/^33^.

### Correlation analysis of gene expression and ageing using self-organizing maps (SOM)

For every gene a tissue-wise Spearman’s rank correlation coefficient was computed based on the expression and age. Similarly, we computed Spearman’s rank correlation coefficients for expression and sex. The resulting correlation matrix was then filtered, such that only genes that were significantly correlated with either age or sex in any tissue (P < 0.01 after false discovery rate adjustment with the Benjamini-Hochberg procedure^32^) were considered. This matrix was then used to create a self-organizing map (SOM)^34^ with the Kohonen R package (version 3.0.8)^35^. In addition to Spearman’s rank correlation coefficients for comparing gene expression and sex we also tested other measures, such as log transformed P-values from one-sided Wilcoxon Ranksum test (testing both, i.e. “greater” and “less” alternatives and flipping the sign if the “greater” alternative yielded a lower P-value) and Somers’ D. Since the different approaches gave similar results we stayed for gender and age with the Spearmann correlation for better comparability.

### Specificity of gene expression for tissues and ages

To identify how specific a gene is expressed at a certain time point or in a certain tissue we employed an approach known from gene set analysis, so-called gene set enrichment analysis (GSEA). Instead of computing how significantly a biological category is enriched in a sorted list of genes, we however computed how enriched a certain tissue, age or pair of tissue/age is enriched in each gene. For each gene we computed 10 enrichment scores for the 10 ages, 17 enrichment scores for the 17 tissues and 170 enrichment scores for the combinations of them. The specificity is defined as the difference between the maximal enrichment and second maximal enrichment for all tissues, time points and combinations. The higher the difference is the more specific is the gene either for a tissue, time point or the combination thereof. Notably, the maximal enrichment score can be translated into p-values, e.g. by random sampling. For better comparability we present however the running sum directly and do not translate them to p-values. Here, the approach is referred to as Sample Set Enrichment Analysis (SSEA).

### Estimating the variance of the data depending on metadata

To estimate the variance in the data depending on age, tissue or gender we made use of Principal Variance Component Analysis (PVCA) as implemented in the Bioconductor Package *pvca*. PVCA combines the strength of principal component analysis and variance components analysis (VCA). Originally it has been applied to quantify batch effects in microarray data. In our case we however do not provide experimental batches but rather groups of meta data as input.

## Code Availability

All code used for analysis will be available upon publication.

## Data Availability

Raw data are available on GEO (GSE132040).

